# “Compressive learning” scaffolds higher-order network structure to enhance human knowledge acquisition

**DOI:** 10.1101/2024.08.19.608587

**Authors:** Xiangjuan Ren, Muzhi Wang, Tingting Qin, Fang Fang, Aming Li, Huan Luo

## Abstract

Humans naturally seek knowledge, yet integrating vast, fragmented information remains challenging. Traditionally, knowledge acquisition has relied on random walks within network—an unguided and inefficient process. We introduce compressive learning, a framework that embeds higher-order structural features—specifically node-degree inhomogeneity—into pre-learning trajectories to scaffold more efficient learning. Across two large-scale experiments, we demonstrate that scale-free networks—due to their pronounced node-degree inhomogeneity—are more compressible and learnable than other network types, and confirm the efficacy of the compressive learning approach. Magnetoencephalography (MEG) recordings reveal that compressive pre-learning enhances structured neural representations in the dorsal anterior cingulate cortex (ACC). A hypergraph-based two-stage model further reveals that compressive learning constructs a network skeleton of hyperedge-defined substructures that more effectively accommodate new inputs. Together, our results highlight the central role of higher-order network structure in human learning and offer a strategic approach to effectively “connect the dots.”

## Introduction

Humans are inherently driven to seek knowledge. Yet in today’s era of information explosion, we often become overwhelmed and lose sight of the bigger picture^1^. Individuals face significant challenges in constructing integrated knowledge frameworks from data that far exceed their processing capacities^2,3^. Understanding the cognitive and computational mechanisms that support knowledge acquisition—and developing efficient strategies to scaffold that process—is therefore critical for individuals and societies alike^4–11^.

Knowledge learning can be conceptualized as navigation through a network, where discrete pieces of information and their relationships are represented as nodes and edges, respectively^12–14^. From this perspective, learning involves uncovering the hidden structure of the network—a process akin to “connecting the dots.” This can be empirically studied using the network learning paradigm^12,15–17^, where participants are exposed to sequences of meaningless images embedded in a predefined network, and each edge dictates the transition probability between images (Figure 1a). As participants predict upcoming images through trial-and-error, improvements in accuracy reflect their acquisition of the underlying network structure.

**Figure 1.**
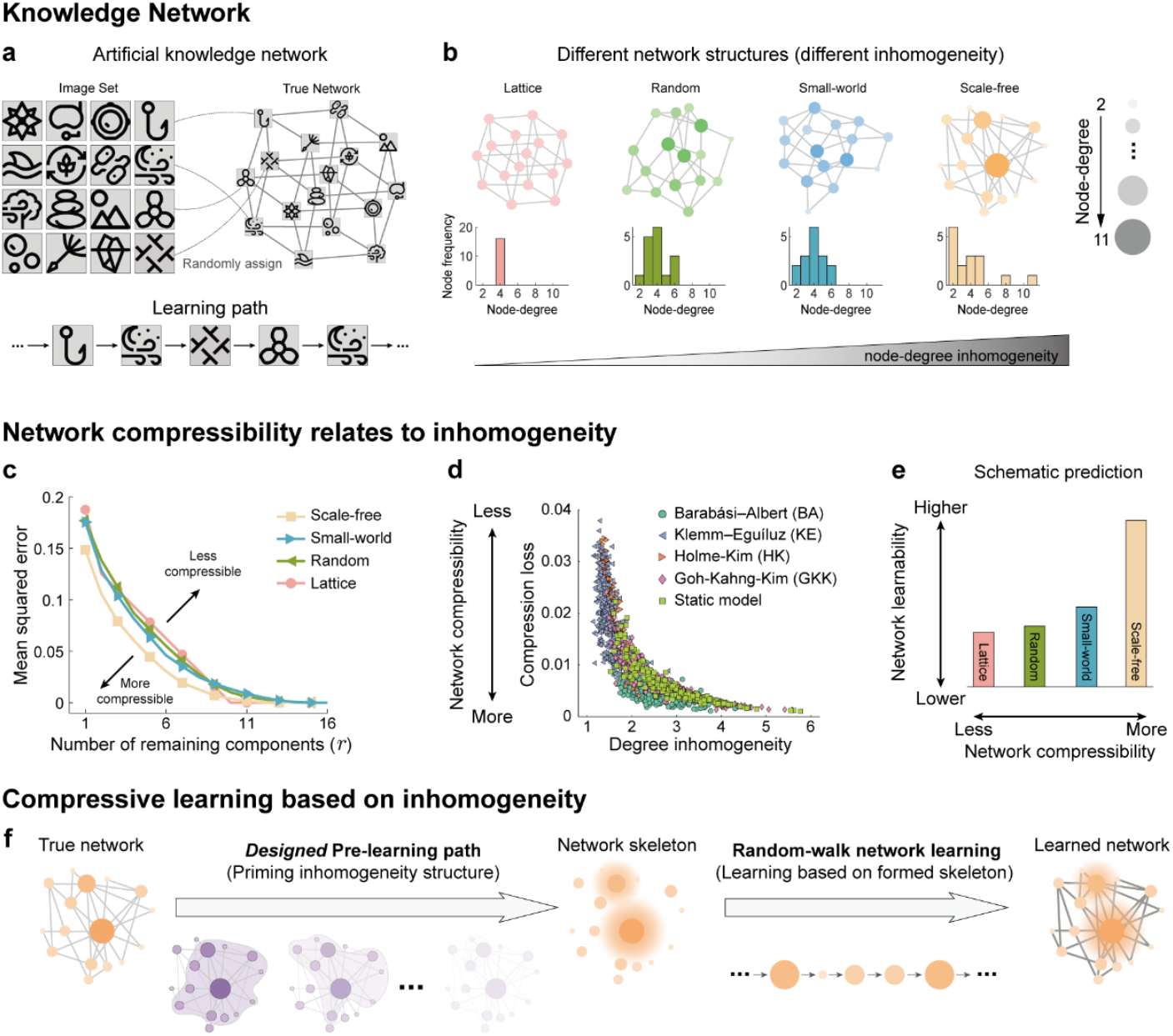
The schematic diagram of compressive learning. **a**, Sixteen images (left) are randomly assigned as nodes in an underlying network (right). Participants were asked to learn the network through a pair-by-pair learning path (bottom). **b**, Four underlying networks with increasing inhomogeneity are designed. Up: Lattice (pink), Random (green), Small-world (blue), Scale-free (orange) networks. Node-degree is denoted by node size and brightness. Bottom: the distribution of node-degree of the four networks. **c**, Compressibility is measured by how less compression loss the compressed network will have after SVD compression. We measure the compression loss with the area under the mean squared error (MSE) curve (see Methods). The smaller the area, the higher compressibility. Scale-free is the most compressible network among the four. **d**, Compressibility increases with node-degree inhomogeneity, indicated by decreasing compression loss. Networks of different inhomogeneity levels are generated from multiple network generation models, denoted by different colors and shapes. **e**, Schematic prediction. As the network compressibility along with inhomogeneity increases, participants learn the network better. **f**, Schematic diagram of compressive learning: Designed pre-learning path primes the inhomogeneous structure of a network, builds a network skeleton, and thus, facilitates the subsequent random-walk learning.

While prior research has shown that abstract structure facilitates the “connecting the dots” process and supports generalization to new inputs^9,12,16,18–21^, the core structural properties that determine a network’s learnability remain unclear. Critically, conventional approaches often use random-walk paths for learning—bottom-up, unguided sampling that is inherently inefficient and slow. Here, we sought to develop a new approach that embeds key structural properties into a pre-learning path to enhance later network learning. This idea also aligns with constructivist theory^22,23^, which posits that learning is an active process where new information is gradually integrated into pre-existing mental models.

Recent work suggests that higher-order network structures—such as community organization and node-degree inhomogeneity—support the mental representation of knowledge networks^14,17,24,25^. These properties are also widespread in real-world systems, further highlighting their role in efficient information storage^26–30^. We focus on node-degree inhomogeneity, a key higher-order property of network structure. As illustrated in Figure 1b, a lattice network exhibits the lowest inhomogeneity, with all nodes having the same number of connections. Random and small-world networks show moderate inhomogeneity, characterized by approximately normal node-degree distributions. In contrast, scale-free networks demonstrate the highest inhomogeneity, following a long-tail distribution in which a few nodes possess a disproportionately large number of connections.

Inspired by the concept of “compressive sensing” in engineering^31,32^, we hypothesize that higher-order structural properties like node-degree inhomogeneity are related to network compressibility—and in turn impact its learnability. More compressible structures are presumably stored more efficiently, enabling more effective network learning. A theoretical compressibility analysis of four network types confirmed that scale-free networks are more compressible than others (Figure 1c), and that greater node-degree inhomogeneity reliably predicts higher compressibility across multiple quantification methods (Figure 1d). We thereby formulated *Hypothesis I*: scale-free network structure, due to its strong inhomogeneity, will exhibit superior learnability in humans compared to the other three network types, even when all networks embed the same image sets (Figure 1e).

Building on this, we proposed *Hypothesis II*, termed “compressive learning” (Figure 1f). We hypothesize that a pre-learning path designed to highlight the inhomogeneous structure of a network can establish a scaffold that facilitates subsequent learning. This approach departs from conventional random-walk network learning by embedding structure-based guidance early in the learning process, thereby acting as a compressive, structure-driven pre-training strategy, rather than a purely bottom-up, unguided process.

To test these ideas, we conducted two large-scale behavioral experiments (*N* > 400), which confirmed both the superior learnability of inhomogeneous (scale-free) networks and the effectiveness of the compressive learning approach. Magnetoencephalography (MEG) recordings further revealed that compressive pre-learning enhances the formation of structured neural representations in the dorsal anterior cingulate cortex (ACC). Finally, we developed a computational model based on hypergraph theory, showing that compressive learning facilitates the formation of a network skeleton composed of hyperedge-defined substructures, which efficiently integrates subsequent inputs. Together, these findings underscore the important role of higher-order network structure in human learning and demonstrate how it can be strategically harnessed to more effectively “connect the dots.”

## Results

### Behavioral paradigm for knowledge network learning

The behavioral task was adapted from previous network learning paradigms^17^. A series of 16-node, undirected and unweighted networks were created in which the 1-step transition probabilities between individual nodes are pre-determined accordingly (Figure 2a). 16 black-and-white images were randomly assigned to the 16 nodes of the network (Figure 2a, left), constituting the to-be-learned artificial knowledge network. Next, image sequences were generated based on a random walk path within the network (Figure 2b, right). Subjects performed continuous prediction through trial-and-error without being told about the actual underlying network. On each trial, subjects were presented with a Cue image and predicted its succeeding image from the Target-Distractor image pair (Figure 2b, left), and the Target image on the current trial would automatically become the Cue image in the next trial (Figure 2b, middle). We monitored subjects’ mouse trajectories during decision-making and performed a trial-wise binary classification analysis to quantify network learning performance (Figures 2c and S2). Specifically, the mouse cursor would be closer to the target than to the distractor if subjects had learnt the transitional relations. Moreover, we employed a “hint system” and a “coin system” to motivate learning (see details in Methods and Figure S1).

**Figure 2.**
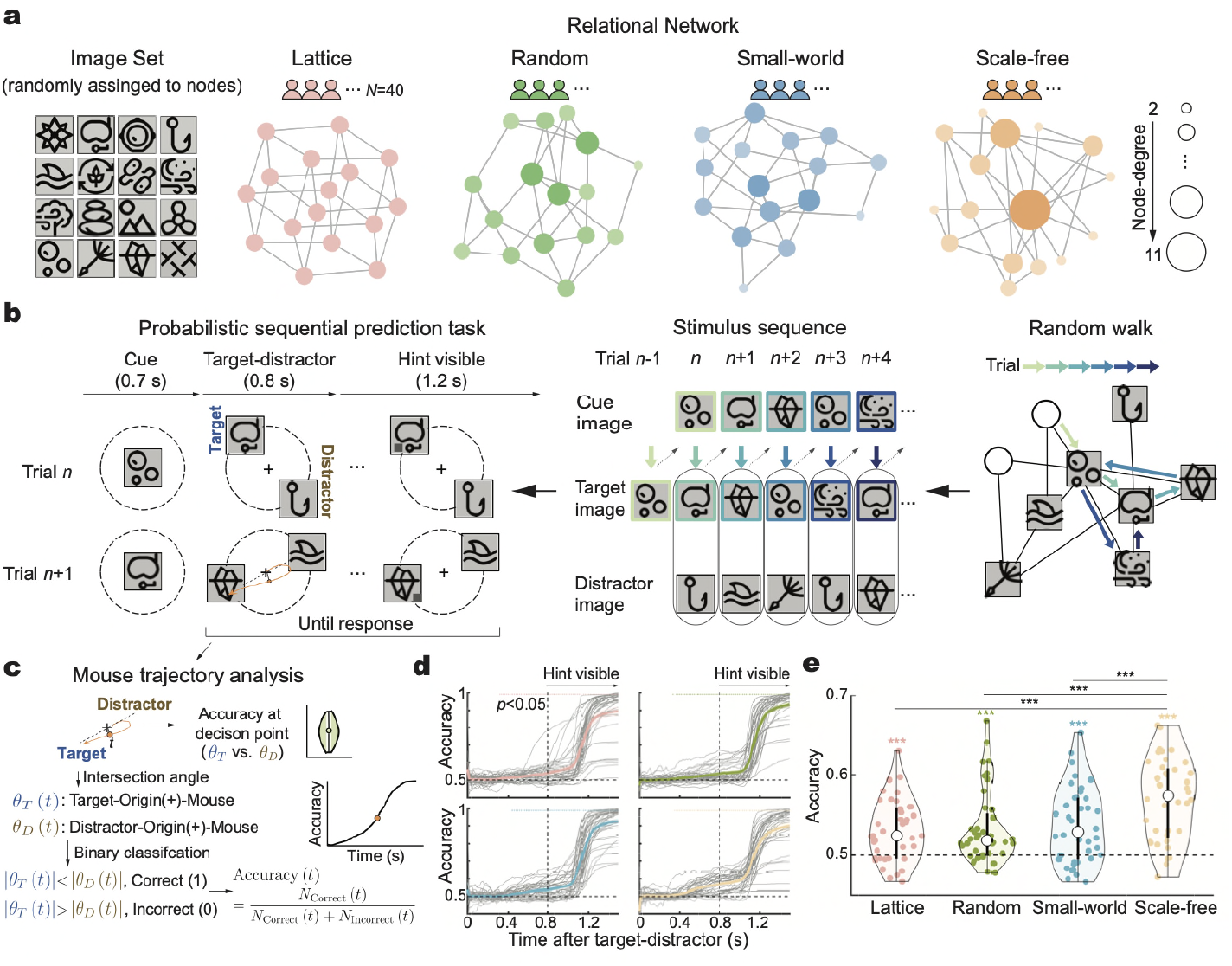
Network learning paradigm and behavioural performance (Experiment 1). **a**, Four groups learned four types of 16-node, undirected and unweighted transitional networks: Lattice (pink), Random (green), Small-world (blue), Scale-free (orange) networks with matched basic properties. Node-degree is denoted by node size and brightness. The same 16 black-and-white images (left) are randomly assigned to the 16 network nodes per subject. **b**, Sequential prediction task. Left: after cue image (“Cue”), subjects predict its succeeding image by mouse-clicking one of the two images (“Target-Distractor”). After 0.8 s, a gray square gradually appears on the target as a hint (“Hint visible”). Response before hint is encouraged and yields higher rewards. Middle: current-trial target image becomes the next-trial cue (dashed arrows). Right: target image sequence is generated via random-walk within corresponding network. Light green to dark blue arrows denote progression across trials. **c**, Time-resolved mouse trajectory analysis. Left: at each time point, angle between mouse cursor position and target or distractor images are computed as *θ*_*T*_(*t*) and *θ*_*D*_(*t*), respectively. Smaller *θ*_*T*_(*t*) than *θ*_*D*_(*t*) indicates correct prediction and larger *θ*_*T*_(*t*) than *θ*_*D*_(*t*) indicates incorrect prediction. Accuracy (*Accuracy*(*t*)) is calculated as the proportion of correct classifications. Right: overall prediction accuracy at decision point. **d**, Time-resolved prediction accuracy of four networks. Gray and colored lines denote individual subjects and grand average. Vertical dotted line indicates hint onset. Colored dots on top indicate group-level significance (compared to 0.5 chance level; one-sample *t*-test, two-sided, *p <* 0.05, *FDR* correction across time). **e**, Overall prediction accuracy of four networks. Colored stars: above-chance accuracy within a group (compared to chance-level 0.5; one-sample *t*-test, two-sided). Black asterisks: post-hoc comparison with Bonferroni correction between groups (GLMM1, ***: *p* < 0.001).

### Inhomogeneity structure facilitates knowledge network learning

Experiment 1 (*N* = 160) directly tested *Hypothesis I* by examining whether a network with high node-degree inhomogeneity would demonstrate superior learnability, given its high compressibility (see Figure 1e). To this end, we constructed four types of 16-node networks (Figure 2a): Lattice (pink), Random (green), Small-world (blue), Scale-free (orange). These networks are matched in the number of nodes (16), edges (32), and average node degree (4), but differ in node-degree inhomogeneity (Figures 2b and S1).

Notably, the Lattice network exhibits the lowest inhomogeneity, with each node connected to exactly four others. The Random and Small-world networks display moderate inhomogeneity with approximately normal node-degree distributions. In contrast, the Scale-free network shows the highest inhomogeneity, characterized by a few highly connected nodes. The same set of 16 images were embedded into each network and shuffled across participants to avoid image-related confounds.

A total of 160 participants were randomly assigned to one of the four network conditions (40 per group). Each group learned its assigned network through image sequences generated by random-walk paths within the respective network (Figure 2a). Participants completed 1,000 trials divided into five blocks. To assess learning performance, we conducted a trial-wise binary classification analysis of mouse trajectories (Figure 2c). As shown in Figure 2d, all groups exhibited above-chance prediction before hints, i.e., moving the mouse toward the target rather than the distractor. Crucially, the Scale-free group demonstrated the highest learning performance (Figures 2e and S3–S5). A generalized linear mixed-effects model (GLMM), which controlled for recency, node degree, and other potential confounders, confirmed the superiority of the Scale-free network (ANOVA from GLMM1 with likelihood ratio tests, main effect of network and trial numbers: χ^2^(3) =143.92, *p* < 0.001, χ^2^(1) =26.71, *p* < 0.001, interaction effect between network and trial numbers: χ^2^(3) = 4.45, *p* = 0.216; post-hoc comparisons with Bonferroni correction from GLMM1, Scale-free vs. Lattice: *z* = 10.63, *p* < 0.001; Scale-free vs. Random: *z* = 9.81, *p* < 0.001; Scale-free vs. Small-world: *z* = 8.68, *p* < 0.001; see the full statistical reports in the Supplementary Information).

In sum, these findings support *Hypothesis I*, suggesting that the compressibility of network structure—reflected in higher-order node-degree inhomogeneity—enhances network learnability in humans.

### Compressive learning promotes knowledge network learning

Experiment 2 (*N* = 240) tested *Hypothesis II* (“compressive learning”) by examining whether emphasizing a network’s inhomogeneous structure during pre-learning could establish a core scaffold that facilitates subsequent random-walk learning (see Figure 1f). The Scale-free network was used, given its pronounced higher-order inhomogeneity, making it ideal for testing this hypothesis.

In a scale-free network, nodes can be ranked by node degree, from low-degree leaf nodes (small circles) to high-degree hub nodes (large circles) (Figure 3a, left). While typical random-walk paths (as used in Experiment 1) visit all nodes probabilistically, we designed pre-learning sequences that explicitly emphasized node-degree structure. Specifically, a 200-image sequence from a random walk (*RandomWalk*) was reordered based on node degree—either in descending (*HubToLeaf*) or ascending (*LeafToHub*) order—yielding three pre-learning paths: *RandomWalk, HubToLeaf*, and *LeafToHub* (Figures 3a and 3b). All three sequences contained the same images with identical frequencies, differing only in order.

**Figure 3.**
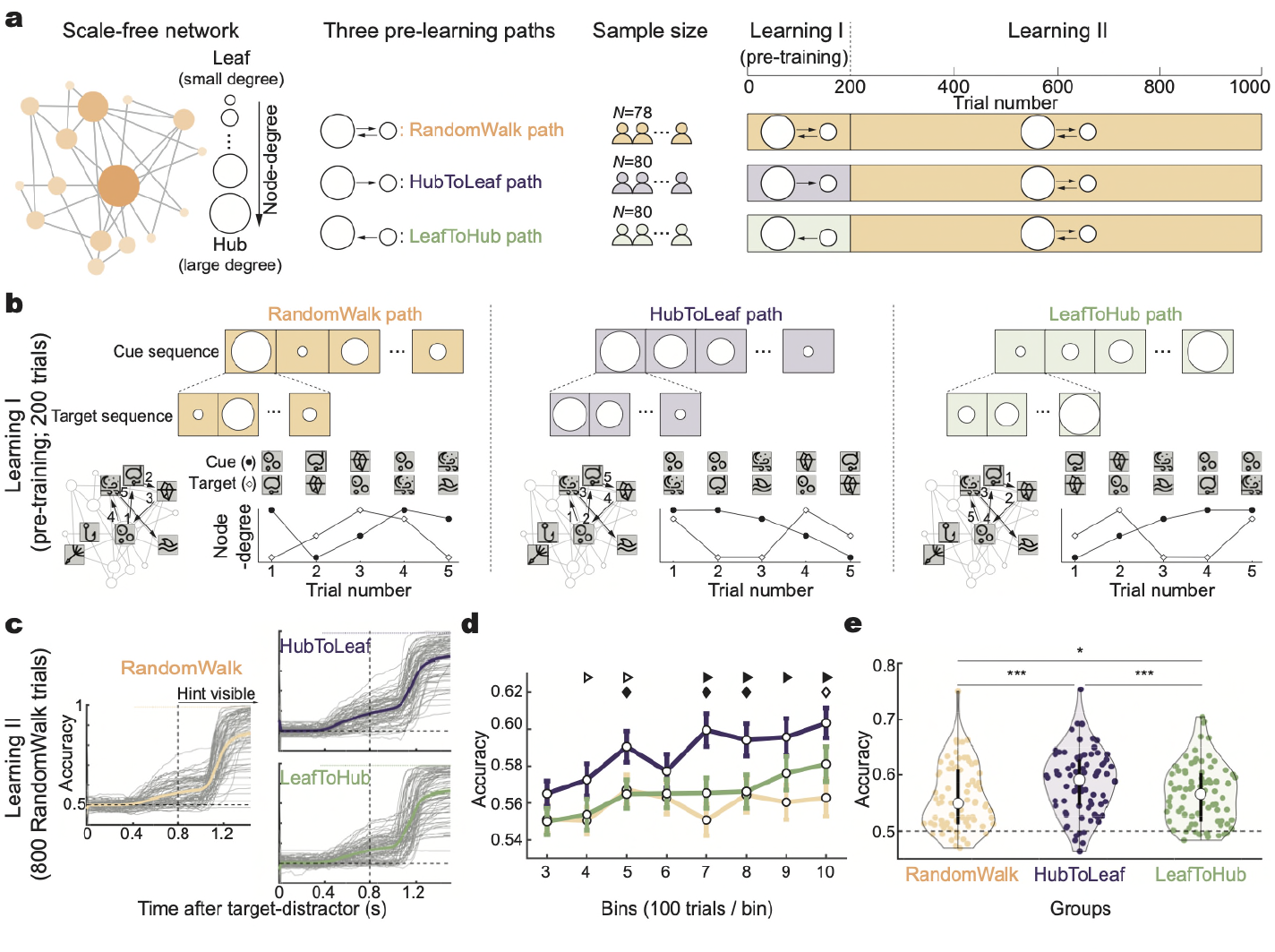
“Compressive learning” facilitates subsequent learning performance (Experiment 2). **a**, Left: inhomogeneity properties of scale-free network, containing Hub (large circle; more connections) and Leaf nodes (small circle; fewer connections). Middle: three pre-learning paths – *RandomWalk* (yellow), *HubToLeaf* (purple), *LeafToHub* (green). Right: during Learning I (200 trials), three groups learned sequences generated by different pre-learning paths (*RandmWalk, HubToLeaf, LeafToHub*). During Learning II (800 trials), they learned identical random-walk image sequences (yellow). Only learning II performance was compared between groups. **b**, Generation of three pre-learning paths (Learning I). Left: *RandomWalk* path is generated by 200-step random walks (arrows with number) visiting hub and leaf nodes in a mixed manner. Middle: HubToLeaf path is the reordering of the 200-length RandomWalk path based on cue and target images’ node-degree in descending order. Right: *LeafToHub* path is the reordering of *RandomWalk* path in ascending order. The three pre-learning paths contain the same images with same frequencies but in different orders. **c**, Time-resolved prediction accuracy for three groups (Learning II). Gray and colored lines denote individual subjects and grand average. Vertical dotted line indicates hint onset. Colored dots on top indicate group-level significance (compared to 0.5; one-sample *t*-test, two-sided, *p <* 0.05, *FDR* correction across time). **d**, Temporal evolution of prediction accuracy for three groups (Learning II). Triangle: *HubToLeaf* vs. *Random*; diamond: *HubToLeaf* vs. *LeafToHub*. Filled, *p <* 0.01; hollow: 0.01 ≤ *p <* 0.05 (GLMM2 and Bonferroni corrected post-hoc comparisons in each time bin; *FDR* correction across bins). **e**, Overall prediction accuracy for three groups (Learning II). *: 0.01 ≤ *p* <0.05; ***: *p <* 0.001 (GLMM2 and post-hoc comparisons on all trials during Learning II).

Specifically, the *HubToLeaf* path introduced relationships between hubs and their neighbors first, followed by progressively lower-degree nodes. In contrast, *LeafToHub* began with leaf-node relationships, introducing hubs last. These reorderings disrupted the probabilistic continuity of the original random-walk path, instead highlighting the underlying inhomogeneity of the network to varying degrees. Based on the centrality of hubs and the primacy effect in learning, we hypothesized that the *HubToLeaf* path would better scaffold the network structure, thereby facilitating subsequent learning.

A total of 238 new participants were randomly assigned to one of the three pre-learning conditions. Each group viewed an image sequence from one of the pre-learning paths (200 trials; Learning I), followed by the same *RandomWalk* sequence used in all groups (800 trials; Learning II) (Figure 3a, right). Critically, only Learning II performance was compared between groups to examine the pre-learning paths’ impacts, and therefore any learning difference would arise from the influence of pre-learning path. As in Experiment 1, learning was assessed using trial-wise classification of mouse trajectories, and all groups performed above chance before receiving hints (Figure 3c). The 800 trials in Learning II were divided into eight bins (100 trials each) to analyze learning progression. As shown in Figure 3d, all groups began with similar performance, but the *HubToLeaf* group gradually surpassed the others and maintained this advantage (see GLMM2.1 in the Supplementary Information). Averaged performance across all Learning II trials followed a similar trend, confirming the superiority of the *HubToLeaf* condition (Figure 3e; ANOVA from GLMM2.2 with likelihood ratio tests, main effect of group and trial numbers: χ^2^(2) =124.68, *p* < 0.001, χ^2^(1) = 22.71, *p* < 0.001, interaction effect between group and trial numbers: χ^2^(2) = 9.35, *p* = 0.009; post-hoc comparisons with Bonferroni correction from GLMM2.2, *HubToLeaf* vs. *RandomWalk*: *z* = 10.72, *p* < 0.001; *HubToLeaf* vs. *LeafToHub*: *z* = 8.03, *p* < 0.001; *RandomWalk* vs. *LeafToHub*: *z* = –2.77, *p* = 0.017). These findings were further replicated in another offline experiment (Figure S6).

In sum, the results support the “compressive learning” (*Hypothesis II*): briefly priming the inhomogeneity network structure (*HubToLeaf* pre-learning path) would facilitate subsequent random-walk network learning.

### Dorsal ACC tracks network structure establishment facilitated by compressive learning

After confirming the effectiveness of “compressive learning”, we next examined the underlying neural mechanism using MEG recordings. A total of 40 new subjects were recruited and assigned randomly to the *HubToLeaf* (purple) or *LeafToHub* (green) groups (20 participants per group). The experimental paradigm mirrored that of Experiment 2 with minor modifications to accommodate MEG constraints (Figures 4a and S1b). During Learning I, participants were exposed to either the *HubToLeaf* or *LeafToHub* image sequence, followed by an identical *RandomWalk* sequence across both groups for Learning II (600 trials). Crucially, only neural activity from Learning II was analyzed and compared between the groups.

**Figure 4.**
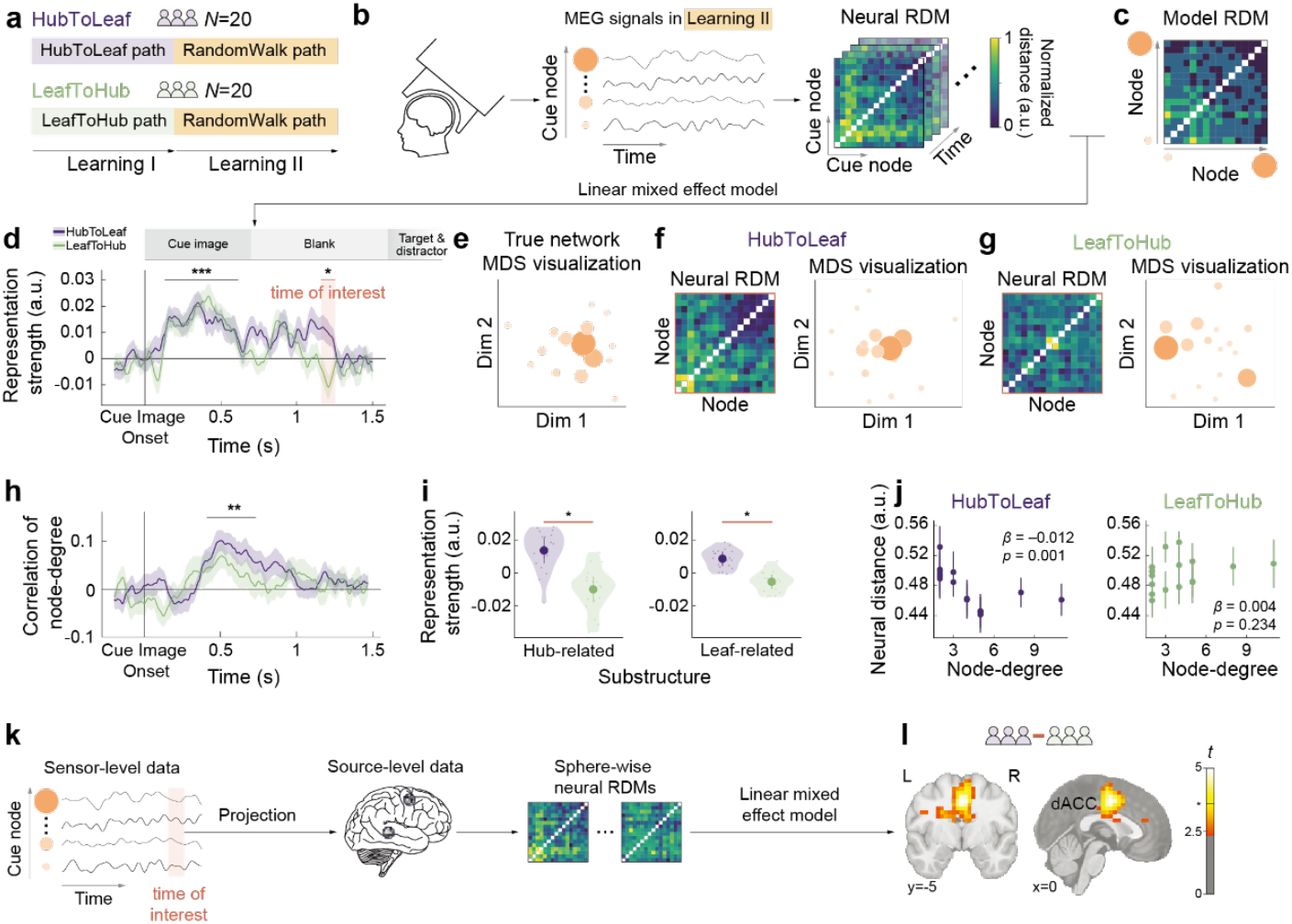
“Compressive learning” enhances network structure formation in dorsal ACC (Experiment 3, MEG). **a**, Two groups participated in Experiment 3 consisting of Learning I (200 trials; *HubToLeaf* or *LeafToHub* paths) and Learning II (600 trials; same *RandomWalk* path). Only Learning II is analyzed. **b**, Each trial is labelled by the identity of cue image within network labels each trial. Multivariate neural dissimilarities are calculated for all node pairs at each time point, resulting in time-resolved neural representational dissimilarity (neural RDM). **c**, Model RDM is constructed based on the corresponding minimal transitional distance within the network. Regressing Neural RDM by model RDM (linear mixed model, LMM1), yielding neural representational strength of network structure. **d**, Time-resolved neural representational strength of network structure during Learning II, for *HubToLeaf* (purple) and *LeafToHub* (green) groups (cluster-based permutation test; ***, *p* < 0.001; *, *p* < 0.05). Time window with significant group differences (shades in orange) is used as the time-of-interest for further analysis. **e**, Two-dimensional representation of the ground-truth network structure (multidimensional scaling of model RDM; MDS). Dots denote images within network. Dot size indexes node-degree. **f, g**, MDS of neural RDM within the time-of-interests for *HubToLeaf* and *LeafToHub* groups. **h**, Time-resolved neural correlation of node-degree (cluster-based permutation test; **, *p* < 0.01). **i**, Neural representational strength of sub-network structures – Hub-related and Leaf-related submatrix – within the time-of-interests, for HubToLeaf (purple) and *LeafToHub* (green) groups. **j**, Average neural distances of each node decrease with node degree in the *HubToLeaf* group (left), but not in the *LeafToHub* group (right). **k**, Source-level analysis. Sensor-level data is projected into source space to perform the same RSA analysis within spheres of 5-mm radius. **l**, Searchlight results for *HubToLeaf*-*LeafToHub* group difference on neural representational strength of network structure (Whole-brain t-value map of group effect, LMM1; Asterisk in the color bar: threshold of *p* < 0.05, FDR corrected).

To probe neural representations of network structure, we conducted representational similarity analysis (RSA) on MEG responses following the presentation of cue images (Figure 4b). We hypothesized that the multivariate dissimilarities between cue-evoked neural patterns (neural RDMs) would reflect the transitional distances between nodes in the underlying network (model RDM, Figure 4c). At each time point, the neural RDMs were regressed against the model RDMs, producing a time-resolved index of network representational strength. Due to random shuffling of image-node mappings across participants, these effects cannot be attributed to image identity. As shown in Figure 4d, both groups demonstrated above-chance neural representations of network structure (black line). Notably, during the blank interval preceding the target/distractor display, the *HubToLeaf* group exhibited significantly stronger network representations than the *LeafToHub* group (LMM1 with cluster-based permutation test, *p* = 0.042).

Further analyses focused on this time window of interest (∼1.2 s post-cue, shaded in orange) confirmed the robustness of this effect. First, multidimensional scaling (MDS) of neural RDMs revealed a hierarchical node organization resembling the ground-truth network (Figure 4e) for the *HubToLeaf* group (Figure 4f), but not for the *LeafToHub* group (Figure 4g). Second, the representational advantage in the *HubToLeaf* group was evident for both hub- and leaf-related substructures (Figure 4i; LMM1, right-sided, hub-related, *β* = 0.012, *t*(44.48) = 2.12, *p* = 0.040; leaf-related, *β* = 0.007, *t*(43.85) = 2.25, *p* = 0.029), indicating a network-wide rather than hub-specific effect. Moreover, both groups showed comparable neural encoding of node-degree information at the early stage but not at the later stage (Figure 4h; cluster-based permutation test, *p* = 0.013; see Figures S8–9 for spatial maps; see Supplementary text for RSA analysis with node degree controlled), suggesting that the enhanced network representations in *HubToLeaf* cannot be attributed to node-degree information. In addition, analysis of neural geometry revealed a hub-attracted distortion in the *HubToLeaf* group (Figure 4j; LMM2, two-sided, *HubToLeaf, β* = –0.012, *t*(4760.00) = –3.27, *p* = 0.001; *LeafToHub, β* = 0.004, *t*(4760.00) = 1.19, *p* = 0.234), consistent with predictions from models like the successor representation^17,26^.

To localize these effects, we projected MEG data from the time window of interest into source space and repeated the RSA. This revealed that the dorsal anterior cingulate cortex (ACC) significantly differentiated the groups in terms of network structure representation (Figure 4kl; LMM1, FDR correction for whole-brain searchlight, peak *p* = 0.004, MNI *X* = 0, *Y* = -5, *Z* = 45). No significant differences were observed in hippocampal regions (Figure S11), and the effect was not explained by differences in univariate activity (Figure S10). Early network representations common to both groups were localized to the bilateral medial frontal cortex and right postcentral cortex (Figure S9, Table S1). Notably, to minimize movement during MEG recording, participants responded via button presses instead of continuous mouse tracking as used in Experiment 2. This less sensitive behavioral measure did not reveal significant performance differences between groups (Figure S7).

In summary, MEG recordings provide convergent neural evidence that compressive learning facilitates the formation of inhomogeneous network structure representations in the human brain. These representations are localized primarily to the dorsal ACC, highlighting its potential role in structuring knowledge based on high-order network properties.

### Hypergraph-based computational model explains compressive learning

Finally, we developed a computational model grounded in hypergraph theory^33,34^ to elucidate the mechanisms of “compressive learning” observed in Experiment 2, which included two stages: Learning I (*HubToLeaf, LeafToHub, RandomWalk* pre-learning; 200 trials) and Learning II (*RandomWalk*; 800 trials). In Learning I, we modeled the formation of an initial network scaffold using hypergraphs, where each hyperedge (*E*_*i*_) comprised a focal node *i* and its directly connected neighbors (Figure 5a and S12). This approach captures not only pairwise connections but also higher-order relationships among sets of nodes. The two pre-learning conditions—*HubToLeaf* and *LeafToHub*— were modeled as distinct sequences of hyperedge construction differing in order and structural emphasis (Figure 5b, left). Specifically, the *HubToLeaf* path (purple) constructs hub-related hyperedges before forming leaf-related ones, whereas the *LeafToHub* path (green) proceeds in the reverse order. Given the primacy effect in learning—where earlier information tends to be more robustly encoded—the *HubToLeaf* condition would create a more structured network scaffold that reflects the inhomogeneous properties of the underlying network. During Learning II, the image sequences generated via random walks were integrated incrementally into the scaffold established in Learning I (Figure 5b, middle). The two-stage model characterizes how new experiences are anchored onto previously formed hyperedges and ultimately yielding the final learned knowledge network (Figure 5b, right).

**Figure 5.**
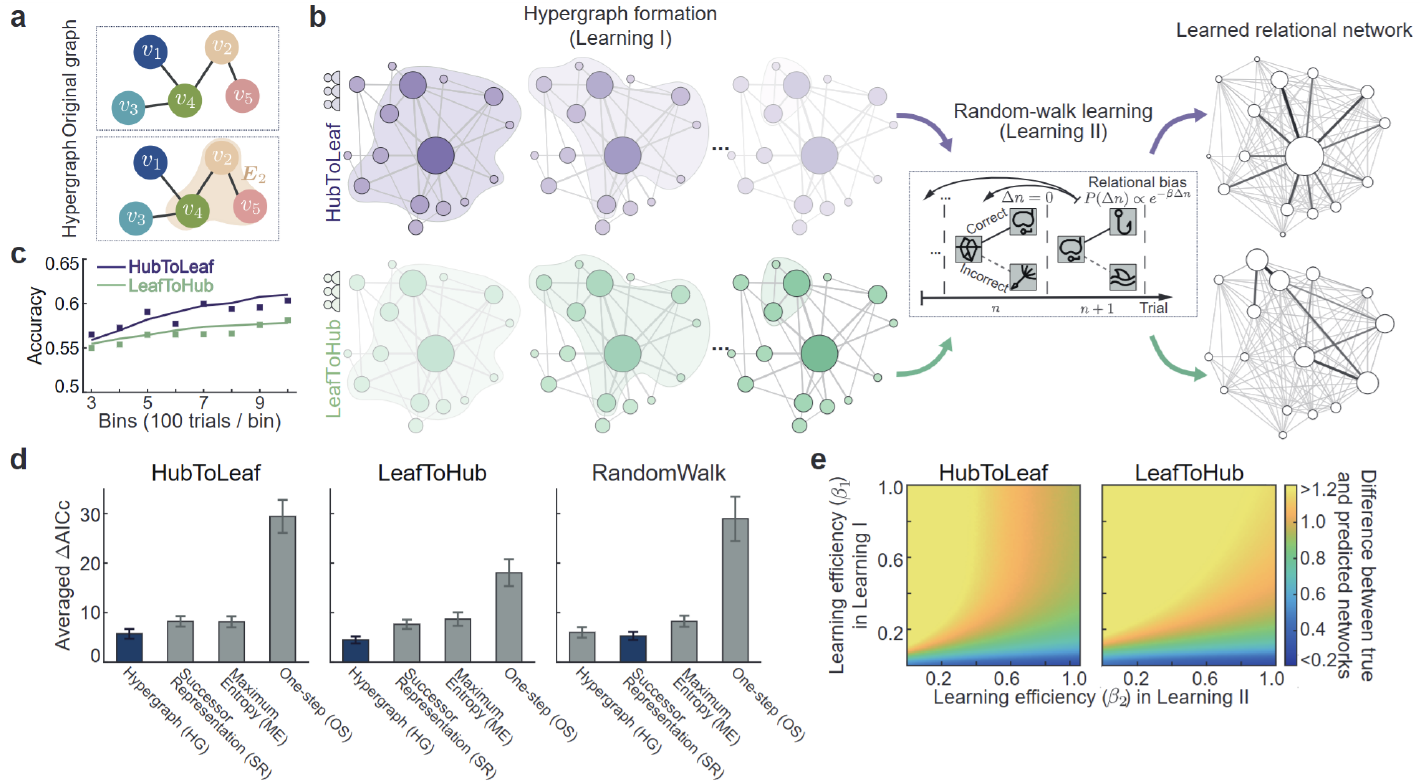
Theoretical model. **a**, Up: Traditional graph theory. Bottom: In the hypergraph, hyperedge *E*_*i*_ consists of node *v*_*i*_ and its all neighbors (shaded plane), e.g., *E*_2_ contains *v*_2_, *v*_4_ and *v*_5_. **b**, Illustration of hypergraph-based “compressive learning” (Experiment 2). During Learning I, for *HubToLeaf* pre-learning path, Hub-related hyperedge is first formed, followed by Leaf-related hyperedge, and for *LeafToHub* group, hyperedges are formed in the reverse order, from Leaf-related to Hub-related hyperedge. In Learning II, subjects would recall previously encountered image sequences, potentially inducing relational biases, captured by the Boltzmann distribution *P*(Δ*n*) = *e*^−*β*Δ*n*^, with Δ*n* denoting the in-between trial interval. Combining these two learning phases yields two learned relational networks, of which the *HubToLeaf*-derived network exhibits higher similarity to the true network than the *LeafToHub*-derived one. **c**, Model fitting (solid line) and behavioral performance (square filled markers) during Learning II. **d**, Model comparisons among Hypergraph (HG), Successor Representation (SR), Maximum Entropy (ME), and One-step (OS) models for *HubToLeaf, LeafToHub*, and *RandomWalk* groups. A lower value of average ΔAIC across subjects indicates a better fit. **e**, Simulated learning performance (KL divergence) between true network and predicted learned network as a function of *β*_1_ (learning I) and *β*_2_(learning II), for HubToLeaf (left) and LeafToHub (right) paths (lower value denotes better performance). Values were truncated to [0.2, 1.2] for better visualization.

We also compared the *hypergraph-based* model against three alternative models that differ in their treatment of higher-order network structure (see Methods). The *one-step learning* model^35,36^ served as a baseline, updating only direct transitions between consecutive items and ignoring higher-order structure. The *maximum entropy* model^37^ captures indirect relationships via probabilistic inference across multiple transitions. The *successor representation* model^26,38,39^, rooted in reinforcement learning, captures extended temporal relationships by predicting future states discounted over time. Notably, none of these models incorporate a hyperedge-based hypothesis.

Model comparisons using the Akaike Information Criterion (AIC) revealed a clear advantage for models incorporating higher-order structure. The *one-step* model performed significantly worse across all conditions, underscoring the importance of higher-order relationships in network learning. Critically, the *hypergraph-based* model not only accounted for the observed *HubToLeaf* vs. *LeafToHub* performance differences (Figure 5c), but also provided the best overall fit among the higher-order models (Figure 5d). Simulations further supported the robustness of the *HubToLeaf* advantage, revealing consistently lower KL divergence for the *HubToLeaf* group across a wide parameter combination of *β*_1_ and *β*_2_ values (Figure 5e and S13).

Taken together, the modeling results reinforce the compressive learning idea: pre-learning paths that emphasize a network’s inhomogeneous structure facilitate the formation of a core structural scaffold, enabling more efficient integration of subsequent information.

## Discussion

We live in an era of information explosion, where data is overwhelming, fragmented, and unstructured—posing a significant challenge to humans’ innate drive for understanding. Conventional random-walk-based network learning operates through a bottom-up, blind sampling process that is inherently inefficient and time-consuming. Here, we introduce a novel approach—compressive learning—which embeds core structural properties (specifically, node-degree inhomogeneity) into a pre-learning trajectory to enhance the efficiency of network learning. This framework is supported by two large-scale behavioral experiments and is further validated through computational modeling and neural evidence. Together, our findings demonstrate that humans are not only sensitive to higher-order structural properties during the process of “connecting the dots,” but can also exploit them more effectively when these properties are presented in a structured and compressive manner that mirrors the organization of the underlying network. These findings have broad implications, with potential applications in fields such as education, cognitive science, and machine learning.

The central concept of compressive learning draws inspiration from compressive sensing^31,32^, where high-dimensional signals are reduced to a few critical components for efficient reconstruction. Analogously, compressive learning decomposes abstract network structures into substructures of varying significance—ranging from hub-related to leaf-related elements—based on node-degree inhomogeneity. Hub-related substructures, with their dense connectivity, provide more global information about the network and are prioritized in the *HubToLeaf* pre-learning path. This ordering is hypothesized to yield a compressible structural skeleton in the brain that facilitates subsequent integration of new information. Conversely, the *LeafToHub* path emphasizes lower-information substructures early on, potentially resulting in a less efficient scaffold. Notably, in our hypergraph-based computational model, hyperedges represent sets of nodes comprising a central node and its neighbors, thereby capturing both pairwise and higher-order relationships. The model simulates the scaffold formation during *Learning I* and the incremental integration of new inputs during *Learning II*. The framework also resonates with constructivist theories in education, which posit that learners actively construct knowledge by integrating new information with pre-existing mental models.

Humans are naturally inclined toward statistical learning^40,41^—the ability to track associations between sequentially presented items over time. Ideally, these local, pairwise associations could coalesce into a coherent network representation. However, local learning is often limited by constraints in memory and attention^3,42,43^, leading to fragile and error-prone associations. In contrast, higher-order network properties—such as community structure and node-degree inhomogeneity—capture non-local regularities that facilitate a more global understanding of the network^14,30^. While such representations may abstract away finer local details^24,44^, they provide an efficient scaffold for integrating new information and generalizing beyond observed data. These higher-order features are also pervasive in real-world networks^27,45–48^, further underscoring their relevance in efficient information encoding. Our theoretical analysis confirms that networks with stronger inhomogeneity are more compressible, suggesting that such structural asymmetries are particularly informative. Crucially, we provide empirical evidence that node-degree inhomogeneity enhances learnability and can be leveraged to design more effective pre-learning sequences.

MEG recordings constitute neural evidence for compressive learning. Early neural representations—common across both *HubToLeaf and LeafToHub* groups—emerge in bilateral superior frontal and right inferior parietal cortices, consistent with prior findings^49^. However, later activity specific to the *HubToLeaf* pre-learning group is observed in the dorsal ACC, prior to decision-making. The ACC is known to support high-level cognitive functions such as hierarchical decision-making^49–52^, foraging^53^, learning^54,55^, context encoding^56^. Our results thus extend its functional repertoire to include the formation and application of structured network representations. Importantly, control analyses confirm that these ACC effects are not attributable to differences in overall activation, node salience, or hub-related substructure. Instead, the ACC appears to track the structural scaffold most consistent with the underlying network topology, highlighting the neural correlates of compressive learning.

We proposed a two-stage model grounded in hypergraph theory to elucidate the computational basis of compressive learning. Model comparisons revealed that frameworks incorporating higher-order relational structure consistently outperformed those based on simpler transition statistics. Among these, our hypergraph-based model provided the best overall fit, likely due to its explicit encoding of inhomogeneity through hyperedges. While future studies are needed to test the model’s generalizability across different network types, the two-stage approach may offer a broadly applicable framework: in *Learning I*, a non-local scaffold is formed from higher-order structural cues, while *Learning II* integrates new experiences onto this foundational skeleton.

From a broader perspective, the two-stage learning framework—involving scaffold formation (Learning I) followed by structure integration (Learning II)—offers a generalizable model for learning from complex, nonlocal input spaces. This process is distinct from curriculum learning^4,57^, which typically utilizes generalization from easy to hard. In contrast, the *HubToLeaf* strategy follows a hard-to-easy trajectory, starting with structurally central but cognitively demanding components (i.e., hubs). Moreover, unlike models that emphasize temporal statistical properties of input sequences^30,37^, our approach explicitly encodes higher-order relationships using hypergraph constructs. Future work could integrate it with traditional machine learning algorithms. Such efforts may uncover fundamental principles of learning from structured data, with potential applications in cognitive science, education, and artificial intelligence.

## Materials and Methods

### Subjects

The study had been approved by the Institutional Review Board of School of Psychological and Cognitive Sciences at Peking University (#2019-02-08). Subjects provided written informed consent in accordance with the Declaration of Helsinki.

**Experiment 1**. For the in-laboratory experiment, 20 human subjects (aged 18-30) of each group were recruited from Peking University. All of them had normal or corrected-to-normal vision. The behavioural experiment lasted for 60 min, and subjects received 55+(0∼30) RMB for their time (basic participation fee + reward). For each group in the online experiment, 20 human subjects were recruited from the Prolific platform. The online experiment was decomposed into two sessions. Online subjects were allowed to participate in the 1st session if they were university students, aged between 18 and 30 years, fluent in English, and had an approval rate higher than 90. In the first session, subjects read the instructions and then completed a short examination to prove that they had fully understood the task. Only those subjects who passed the test within 5 attempts in the 1^st^ session were invited to the 2^nd^ session (the main experiment). The average completion time for the 1^st^ and 2^nd^ sessions was ∼10 and ∼50 min separately. In addition to the basic participation fee (£7.25), subjects had the opportunity to obtain an additional bonus (max. £4), which was accumulated across trials depending on their accuracy and RTs.

**Experiment 2**. 78, 80, and 80 subjects from the Prolific platform were recruited for the RandomWalk, HubToLeaf, and LeafToHub groups in Experiment 2, respectively. Subjects who had participated in Experiment 1 were disqualified. For the in-laboratory experiment, 21 human subjects (aged 18-30) for each of the path groups were recruited from Peking University.

**Experiment 3**. 40 subjects (aged 18-30) were recruited from Peking University for the MEG experiment.

### Apparatus and stimuli

In-laboratory subjects were seated approximately 86 cm in front of a 21.5-inch iMac monitor (47.9×26.9 cm, 2,048×1,152 pixels, 60-Hz refresh rate). The display of stimuli and the recording of responses were controlled by the iMac computer using MATLAB R2017a and PsychToolbox-3^58,59^. The visual angle of each image was 2.25°, and all images in a display were equidistantly distributed on an invisible circle that subtended a visual angle of 9°. For the online experiment, the stimuli were presented in the unit ‘height’, defined as a relative measure to the height of the window. Thus, the physical sizes and locations of the stimuli would change with different monitors. The stimuli were 16 cartoon images with abstract semantics (https://www.flaticon.com/packs).

### Transition network

Unbeknownst to the subjects, we used four different transition networks (Figure 2a) in Experiment 1 to generate the image sequences across trials, with each node denoting one image in the image set (Figure 2a, left). The four networks were matched in their basic topological features, such as the same number of nodes (16) and edges (32), and an average node-degree (number of edges that are directly connected to the node) of 4. However, their node-degree distributions differed, resulting in diverse heterogeneity (Figure 1a). The Lattice network was a spherical shape, with each node fully connected to its adjacent four nodes. The Random network was generated based on the Erdős-Rényi model by randomly connecting each pair of nodes with the probability *p*, and the corresponding average degree of a random network was given by *p*(*N* − 1), where *N* is the total number of nodes^60^. The Small-world network was constructed by randomly rewiring the edges on regular (or Lattice) networks with the probability *p* = 0.3^61^. In the current setting, the node-degree distributions of Random and Small-world networks were similar to quasi-normal. The Scale-free network was constructed with the Barabási-Albert model^62^, which possessed a unique feature of extremely high connectivity at very few nodes (the “hubs”). All subjects experienced the same 16 images and the same transition network within a group, but the correspondence between nodes and images was shuffled across subjects. Based on the objective transition network, four corresponding minimal distance matrices (MDM) were constructed, with each cell denoting the minimal transition steps between two nodes.

### Probabilistic sequential prediction task

Each trial started with a cue image, followed by the presentation of target and distractor images (“Image onset”). The target image was directly connected to the cue image in the transition network (one-step transition), while the distractor images were more than one-step transitions from the cue image. In each trial, only one target image and one distractor image were displayed. Subjects’ task was to click the target image using the mouse as soon as possible after the image onset without sacrifice of accuracy, during which their time-resolved trajectories were recorded (Figure 2b).

### Experimental procedure

Each trial started with the cue image on the screen for 0.7 s (except for the 1^st^ trial of each block, which was presented for 2 s), after which one target and one distractor image were presented for 2 s (“Image onset”). The subjects’ task was to select the target image as accurately and quickly as possible through learning and feedback.

To facilitate the network learning, after 0.8 s presentations of alternative images, a small gray square (the ‘hint’) appeared gradually on the target image to indicate the correct answer. Note that subjects were encouraged to make a response before hint onset if they were confident. Once subjects made responses, auditory feedback was presented indicating whether the response was correct, incorrect, or time-out. The feedback screen lasted for 0.7 s.

Subjects received rewards on each trial based on their behavioral performances. Specifically, a correct response before hint would yield 100 virtual coins. Otherwise, the accessible coins decreased linearly with time, declining from 100 to 0 within 0.5 s and from 0 to -50 for the remaining 0.7 s. In other words, before- and after-hint responses yielded higher and lower rewards, respectively. Initially, subjects must rely on the hint, which appeared on the target image with the same gray scale as the background at 0.8 s and gradually increased to the largest gray scale within the next 0.8 s, to make a correct response. Otherwise, time-out or incorrect selection would lead to 50 coins loss. In the laboratory experiment, in order to control the mouse’s moving speed, subjects were informed that the cursor’s position would be forced back to the screen center if their moving speeds were too fast.

### Generation of priming scenarios in Experiments 2 and 3

In Experiment 1, the target image in each trial served as the cue image in the next trial for subjects to make predictions, with the target image sequences being prescribed by a random walk on the transition network. A random walk is a path across a transition network created by taking repeated random steps between any two directly connected nodes^28^. In contrast, in Experiments 2 and 3, the continuity of the cue between target images across trials was broken for the first 200 trials. Specifically, the cue-target-distractor combination in each trial was ranked primarily based on the node-degree of cue image, and then ranked again based on the node-degree of target image under the same cue image (Figures 1f and 3b). Hence, the first 200 trials were sorted in descending order based on the node-degree of both the cue and target images for the HubToLeaf group. Conversely, in the LeafToHub group, the trials were sorted in ascending order. In summary, the three groups (Random, HubToLeaf, LeafToHub) differed only in their trial organisations in the first 200 trials (“priming scenarios”), while the stimulus exposure in the remaining 800 (Experiment 2) or 600 trials (Experiment 3) was similar. We therefore referred to the first 200 trials as the Learning I phase, and the remaining trials with random walk transitions as the Learning II phase. Subjects were not informed about the priming scenarios; they were only told that the cue image would be repeated over a period.

### Time-resolved mouse trajectory analysis

In the laboratory experiment, we recorded the mouse trajectory at a sampling rate of 60 Hz during each trial for each subject. The trajectory initiating from the center point and ending at the target or distractor image could reflect subjects’ internal decision process. For the online experiment, we adjusted the sampling rate to 60 Hz by interpolation due to the different operating systems used by the subjects. At each sampling point within a trial, we calculated the absolute angle between the cursor dwell position (pixel unit for the in-laboratory experiment and height unit for the online experiment) and each alternative image (target or distractor). A smaller absolute angle indicates a stronger decision preference towards a specific image.

We calculated the intersection angle between each sample point of the mouse trajectory and the center of the target or distractor images within a trial, with the center of the screen as the origin. The resulting mouse-origin-target angle (*θ*_*t*_) and mouse-origin-distractor angle (*θ*_*d*_) were then imported into a binary classifier to determine subjects’ internal decision at the current sample point (Figure 2c). Specifically, a correct choice was exported if the mouse’s dwell position was closer to the target, i.e., |*θ*_*t*_ |<|*θ*_*d*_ |; otherwise, an incorrect response was output. By doing this, we transformed a continuous circular variable ([-π, π]) into discrete categorical variable (0 or 1). The accuracy at each sampling point was calculated as the proportion of trials classified into correct (choosing the target). To quantify the overall learning effect, we extracted one classification result from each trial. For the response-before-hint trials (Figure S3, top panel), the classification result at the point when subjects clicked the target or distractor was extracted. For the response-after-hint trials (Figure S3, bottom panel), the landing position right before the hint onset (0.8 s) was used to make a classification.

### MEG Experiment

#### Apparatus and MEG acquisition

Visual stimuli were presented with a projector (Panasonic PT-DS12KE) at a refresh rate of 60Hz. Auditory stimuli were presented with a pair of earphones. Subjects were seated in a magnetically shielded room and reacted with a MEG-compatible response box. Neuromagnetic signals were recorded at 1,000 Hz by a 306-channel system (Elekta-Neuromag) with 102 magnetometers and 204 gradiometers while subjects were seated 1 m in front of a projection screen and performing the task. MRI T1 data were acquired after MEG recording, using an MP-RAGE sequence on a Siemens 3T TRIO scanner, with voxel resolution of 1 × 1 × 1 mm^3^ on a 176 × 192 × 192 grid (echo time = 2.98 ms, inversion time = 1,100 ms, repetition time = 2,530 ms). Eye movement was monitored online by an eye tracker (Eyelink 1000 plus, remote mount) at 1000 Hz. Five-point calibration was performed every 100 trials.

#### MEG preprocessing

The preprocessing of MEG data was implemented with MNE-Python (version 1.3.1)^63^, following the FLUX pipeline^64^. Maxwell filter and Signal-space separation were performed first to remove outside-helmet artifacts. The data were then high-pass filtered at 0.5 Hz and low-pass filtered at 40 Hz, followed by downsampling to 250 Hz. Independent component analysis was then performed on the filtered data. Ocular and cardiac components were manually identified for each subject and then subtracted from the data. All epochs were aligned at the onset of cue image. Only trials with correct responses were included for the following MEG analysis.

#### Representational similarity analysis (RSA)

The general principle of RSA is comparing the representational similarity between the neural activity and a proposed model^65^.

The neural representation dissimilarity matrix (RDM) characterizes the brain’s responses. The neural RDM at time point *t, N*(*t*), is a 16×16 matrix. Each element of the neural RDM measures the dissimilarity of the neural representation between the node images. To obtain a robust estimate of the neural RDM, we performed a resampling procedure to balance the number of trials with different node images and repeated the resampling procedure 100 times. For each iteration, we randomly drew 50 trials with replacement from trials with the same cue image. The MEG data of the 50 trials were then averaged. We performed such drawing and averaging on the MEG data of each cue node. To obtain time-resolved neural RDMs, we calculated the Euclidean distance (1 - Pearson’s *r*) between the topographic maps of different nodes at each time point. Then, we averaged RDMs from the 100 iterations. To de-noise the RDM, the RDM time series were subtracted with the average RDM over a baseline window (from the onset of cue image to 500 ms before the onset).

The model dissimilarity matrix (MDM) *M*, a 16×16 matrix, characterizes the underlying network structure^36^. Each element of the MDM denotes the step of the shortest path between two nodes according to the network graph.

To examine how the representation of network structure changes over time, we analyzed the relationship between neural RDM and MDM at each time point. LMM1 were specified to test whether the subjects represent the network structure and whether group difference exists in representation strength. Cluster-based permutation test was used for correction (see Supplementary Text for details).

For visualization, neural RDMs and MDM are normalized and then projected to 2-dimensional Euclidean space with multidimensional scaling. Multidimensional scaling tries to reconstruct the coordinates of all nodes on a Euclidean map that minimize the summed absolute difference between the distance in dissimilarity matrices and the reconstructed distance. Multidimensional scaling is implemented with Python package *sklearn*^66^.

#### Leaf-related and hub-related submatrices

In the Scale-free network, nodes with more than 7 connections are defined as hub nodes, and the rest nodes are defined as leaf nodes. In Experiment 3, to examine whether the specified path influences the entire network, we fitted LMM1 separately on hub-related and leaf-related submatrices. Both submatrices are subsets of a full dissimilarity matrix. Leaf-related submatrix contains only distances between leaf nodes. Hub-related submatrix is the complement set of leaf-related submatrix, which contains hub distances to both hub and leaf nodes.

#### Source-level RSA searchlight

Sensor-level signals within the two effective time windows (50-ms width, centred at 370 ms and 1,196 ms) were projected to an average brain, respectively. Voxels were evenly spaced 5 mm apart. The same RSA with minor modifications was then performed with searchlight at the radius of 10 mm (see Supplementary Text for details).

#### Node-degree regression analysis

At each time point and each channel, MEG data were baselined and then regressed with node-degree of cue image.

### Statistical Analysis

#### Linear mixed models (LMMs) and generalized linear mixed models (GLMMs)

LMMs and GLMMs were implemented for group-level inference to account for both group-level and participant-level variance and to control nuisance variables. Both variables of interest and variables for control are specified as fixed effects. Specification of random effects followed the “keep-it-maximal” principle to prevent false positive results^67,68^. Random effects and their correlations would be reduced gradually in case of singularity. Recency of nodes, node-degree of target nodes, and minimal distance of distractor nodes are controlled as fixed and random effects. While LMM assumes Gaussian distribution for noise, GLMM assumes binomial response noise. The models are fitted with maximum likelihood estimation. The significance of each fixed effect is determined by Welch-Satterthwaite *t* test for LMM and likelihood ratio test for GLMM. The Bonferroni correction is used in post-hoc comparison. Cluster-based permutation test is used for test in the RSA time course^69^.

#### Cluster-based permutation test for multiple comparison correction

The massive statistical test at each time point might cause excessive type I error. We performed a cluster-based permutation test to identify significant temporal clusters^69^. After shuffling the elements in the MDM *M* and subjects’ group labels, we fitted the LMM1 to shuffled data and performed Satterthwaite *t* test at each time point. Continuous significant time points were selected as clusters (threshold *p* < 0.05), and the cluster with the largest summed *t* was retained. We repeated the permutation for 1000 times to obtain a distribution of summed *t*. Null hypothesis was rejected if any cluster from the unshuffled data was larger than 95% of summed *t* in the surrogate distribution.

In node-degree analysis, since channel dimension was not collapsed as in RSA, significant cluster was determined jointly by both temporal and channel dimension.

#### Network heterogeneity analysis

Typically, the degree heterogeneity^70^ of a network is defined as

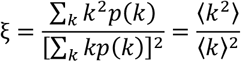

where *p*(*k*) is the distribution probability of degree *k* in the network.

#### Network compressibility analysis

For many natural or technological systems, they are often characterized by the low-rank nature. As demonstrated in ref. 71, using singular-valued decomposition (SVD), the low-rank characteristic of Drosophila melanogaster’s hemibrain connectome can be experimentally verified. According to the Schmidt–Eckart–Young–Mirsky theorem^72^, SVD provides the optimal low-rank approximation of a matrix in terms of the Frobenius norm. Therefore, we can use SVD to measure how the network structure is captured by a low-dimensional structure. For any adjacency matrix *A* ∈ *R*^*n*×*n*^, we can represent it as

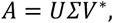

where * denotes the complex conjugate transpose and *U, V* ∈ *C*^*n*×*n*^ are unitary matrices whose conjugate transpose is equal to its inverse, namely *U*^*^*U* = *V*^*^*V* = *I*, where *I* is the identity matrix. The matrix *Σ* = *diag*(*σ*_1_, …, *σ*_*n*_) ∈ *R*^*n*×*n*^ is a diagonal matrix of singular values *σ*_1_ ≥ *σ*_2_ ≥ … ≥ *σ*_*n*_ ≥ 0. In that case, matrix *A* can also be given by the sum of rank-one matrices, weighted by singular values

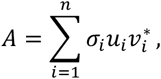

where *u*_*i*_ and *v*_*i*_ are the *i*th column vector of matrices *U* and *V*, respectively. In light of this, SVD offers a hierarchy of low-rank approximations by truncating the sum to include only the leading singular values. Correspondingly, if a matrix is reconstructed more accurately under the same rank approximation, it is considered to be more compressible. We quantify the compressibility of a network using the reconstruction error, here defined as the mean squared error (MSE) between the adjacency matrix of the network and its low-rank approximation across all possible truncation levels, given by

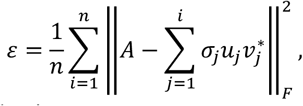

where ‖·‖_*F*_ denotes the Frobenius norm.

#### Theoretical model

Here we utilize a graph to conceptualize the connections (edges) between images (nodes) and represent how the images relate to each other by the adjacency matrix *A* = [*a*_*ij*_] ∈ {0,1}^*s*×*s*^, where *s* is the number of nodes. If images *i* and *j* have a connection, *a*_*ij*_ = *a*_*ji*_ = 1 ; otherwise, *a*_*ij*_ = *a*_*ji*_ = 0. Define the degree matrix *D* = *diag*(*d*_1_, …, *d*_*s*_), where 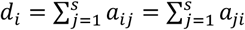.

In Experiment 1, image sequences are generated based on a random-walk path on the underlying network with the corresponding transition probability matrix obtained by *T* = *D*^−1^*A*. Following the work of Lynn et al.^37^, it suggests that the relational biases are inevitable for human subjects (Figure 5b, middle). Consider a sequence of stimuli {*x*_1_, *x*_2_, …, *x*_*N*_, *x*_*N*+1_}, where *x*_*i*_ ∈ {1,2, …, *s*} refers to the image label. At the end of *N* trials, subjects’ biased count of edge (*i, j*) can be estimated by

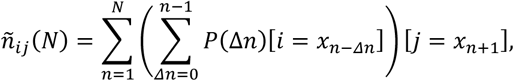

where 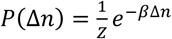 represents the distribution of the relational bias with *Z* serving as a normalization constant such that ∑_Δ*n*_ *P*(Δ*n*) = 1, and [·] is an indicator function, which takes a value of 1 if the equation within the bracket holds and 0 otherwise. The free parameter *β* indicates the learning efficiency of the human subjects. As *N* → ∞, the relative connection weight of edge (*i, j*) can be obtained by dividing both sides by *N*, yielding that

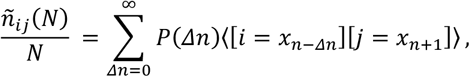

where ⟨·⟩denotes the expectation. For the random walk on the graph,

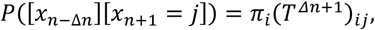

where 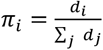 means the stationary distribution of node *i* appearing in the random walk. It then follows that â_*ij*_ = ∑_Δ*n*_ *P*(Δ*n*)*π*_*i*_(*T*^Δ*n*+1^)_*ij*_, where â_*ij*_ indicates the learned internal representation for the edge (*i, j*). We express this in a matrix form, i.e., Â = ∑_Δ*n*_ *P*(Δ*n*)*ΠT*^Δ*n*+1^, where *Π* is a diagonal matrix with its *i*th entry *π*_*i*_.

To convert subjects’ internal beliefs into choice probabilities, we adopt the binomial logistic regression to predict the likelihood of selecting image *i* followed by image *j*, not image *k*

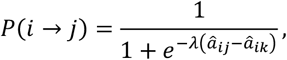

where λ is a free parameter for the inverse temperature to capture choice randomness.

In Learning I of Experiment 2, the stimulus sequence establishes a cohesive learning unit that maps out the connection relation between a node and its neighbours. Human subjects, through these units, form the coarse-grained representation of the underlying network, beyond the dyadic network learning. We rely on the framework of hypergraphs, a potent tool in network science, to capture the process by modeling the basic units as hyperedges and constructing the weighted adjacency matrix to enable the extraction of higher-order network features (Figure 5a). Further, the coupling strength between nodes *i* and *j* in a hyperedge is not merely a binary value but depends on the involved dynamics^73^. We consider that the coupling strength between nodes in a hyperedge is relevant to the importance of nodes, which is intuitively described here in terms of the frequency of occurrence during the compressive learning process. Since normalization is influenced by the relative, rather than the absolute value of the occurrence frequency, we denote the coupling strength between nodes *i* and *j* by *K*_*ij*_ = ∑_*α*_ *k*_*iα*_*e*_*iα*_*k*_*jα*_*e*_*jα*_, where *e*_*iα*_ ∈ {0,1} indicates whether node *i* is in the hyperedge *E*_*α*_, and *k*_*iα*_ = *d*_*i*_ if *i* = *α*, otherwise 1. Therefore, we can derive the newly formulated incidence matrix with 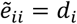 and otherwise 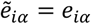. The adjacency matrix of the hypergraph can be represented by 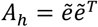 (Figure S13). The above is given the higher-order characterization without considering the effect of local stimulus order on the learned network structure. Due to mental error and resource constraints, a node with a preceding stimulus order has fewer confounders and higher quality in learning its local structure, a phenomenon termed “primary effect” in cognitive science. We specify that the stimulus order of nodes 1, …, *s* is represented by *o*_1_, *o*_2_, …, *o*_*s*_, and *Ω*_*i*_ = {*j*: *o*_*j*_ < *o*_*i*_} denotes the set of nodes before node *i* in the order of stimuli. Further, we assume that

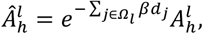

where 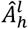 and 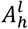 indicate the learned local connection structure of node *l* when the local stimulus order is considered and not considered, respectively. Moreover, the diagonal elements are purposefully set to zero because the network topology does not contain self-edges.

Subsequently, in Learning II, the global features of the network structure can be observed by random-walk stimuli. Given the mixture effect of two learning phases, we define the parameter 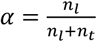 to account for the modulation in the learning process, where *n*_*l*_ denotes the number of trials of Learning I and *n*_*t*_ denotes the number of trials of Learning II. It allows us to derive an analytic expression for the learned network

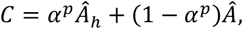

where the parameter *p* is used to measure the degree of nonlinearity in the modulation. Owing to factors such as task difficulty, learning efficiencies may not necessarily be the same in two learning phases. Thus, in order to differentiate, we use *β*_1_ to indicate the learning efficiency for Learning I and *β*_2_ for Learning II.

### Alternative models

#### One-step model

The one-step model assumes that participants learn only the directly experienced one-step transition between two states, with the following counting rule:

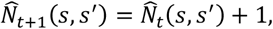

where *s* and *s*^′^ represent the cue image and target image in trial *t*, and 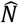 is the internal representation of visiting numbers between states. The transition relationship matrix ***M*** is then derived by normalising these counts, such that for any pair of states *s* and *s*^′^,

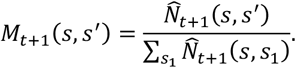

#### Maximum Entropy Model

Rather than focusing solely on direct, one-step transitions, a natural and practical extension incorporates the counts between states that are multi-step away, with each weight captured by the memory-bias probability *P*(Δ*n*), where Δ*n* denotes the step distance in the stimulus sequence. By balancing accuracy and computational complexity in this way, one arrives at a distribution that satisfies the maximum entropy principle—the Boltzmann distribution (*P*(Δ*n*) ∝ *e*^−*β*Δ*n*^), where *β* indicates the learning efficiency. Formally, for the target state *s*^′^, the counting rule obeys

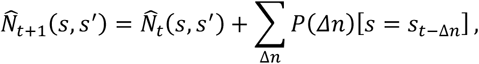

where *s*_*t*_ denotes the state transitioned from the initial state after *t* time-steps. As in the one-step model, the transition matrix is obtained by normalising these modified counts.

#### SR model

In contrast, the SR model assumes a long-term transition relationship between a given state and all its successor states. These relationships are captured in an *n* × *n* matrix, where *n* denotes the 16 states (or nodes) in the experiment. Each cell in the SR matrix represents the expected discounted number of visits from one state (*s*) to another state (*s*^′^), defined as:

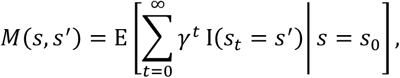

where *γ* is a discount factor (0 < *γ* < 1), and I(·) is a one-hot vector with a value of 1 for the successor state *s*^′^.

The SR model was learned and updated on a trial-by-trial basis using the temporal difference learning^35^:

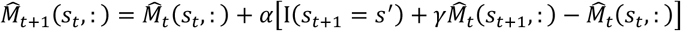

where 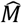 is the internal representation of the transition matrix updated through SR model; *α* is the learning rate (0 < *α* < 1). Specifically, for any transitions from *s*_*t*_ in trial *t* to *s*_*t*+1_ in trial *t* + 1, the successor matrix 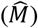 would be updated based on the temporal difference error in the square bracket.

#### Model fitting and comparison

For each subject, we fit the models to subjects’ choice classified through mouse trajectory in each trial using maximum likelihood estimates (MLE). Given the choice in each trial has a choice probability, *p*(*C*_*t*_), the likelihood function derived from binomial distribution was used to describe the relationship between subjects’ responses and the model predictions. The *fminsearchbnd* (J. D’Errico), a function based on *fminsearch* in MATLAB (MathWorks), was used to search for the parameters that minimized negative log likelihood. To verify that we had found the global minimum, we repeated the searching process for 100 times with different starting points. We compared the goodness of fit of each model based on the Akaike information criterion with a correction for sample sizes (AICc)^74,75^. Note that the AICc values punishes number of free parameters, and lower AICc values indicate better fitting performance. Specifically, in each subject, the model with lowest AICc was used as a reference to compute ΔAICc for all models.

## Acknowledgements

This work was supported by the Science Fund for Creative Research Groups of the National Natural Science Foundation of China (T2421004 to F.F., A.L. and H.L.), the National Science and Technology Innovation STI2030-Major Project 2021ZD0204100 (2021ZD0204103 to H.L.), National Key Research and Development Program of China (2022YFA1008400 to A.L.), National Natural Science Foundation of China (31930052 to H.L., 62173004 to A.L.). We thank the Center for MRI Research at Peking University in Beijing, China, for assistance with data acquisition. We also thank Prof. Nai Ding and Prof. Jian Li for their comments on the manuscript and Dr. Haoyang Lu for statistical consultation.

## Author contributions

X.R., A.L., and H.L. originally conceived the study. X.R. and M.W. performed the experiments. X.R., M.W., and T.Q. analysed the data. T.Q., A.L., and H.L. performed theoretical analyses. X.R., M.W., T.Q., F.F., A.L., and H.L. wrote the paper.

## Data and materials availability

All data and codes will be made publicly available upon acceptance of the paper.

